# S^2^-PepAnalyst: A Web Tool for Predicting Plant Small Signalling Peptides

**DOI:** 10.1101/2024.08.02.606319

**Authors:** Kelly L. Vomo-Donfack, Mariem Abaach, Ana M. Luna, Grégory Ginot, Verónica G. Doblas, Ian Morilla

**Affiliations:** Université Sorbonne Paris Nord, LAGA, CNRS, UMR 7539, F-93430, Villetaneuse, France; University of Malaga, Department of Applied Mathematics, MLiMO, 29010, Málaga, Spain; MAP5 UMR CNRS 8145, Université de Paris, 45 rue des Saints-Pères, 75006 Paris, France; Instituto de Hortofruticultura Subtropical y Mediterránea La Mayora (IHSM), Universidad de Málaga-Consejo Superior de Investigaciones Científicas, 29010, Málaga, Spain

**Keywords:** Small signalling peptides, Plant signal peptide prediction, Machine learning in plant biology, Plant development and stress response

## Abstract

Small signalling peptides mediate cell-to-cell communication playing essential roles in plant growth, development, and stress responses. They specifically bind to the extracellular domain of receptors to trigger biochemical and physiological responses. Despite their significance, accurately identifying novel signalling peptides remains challenging due to their structural diversity, low abundance, and highly specific expression patterns. Here, we present S²-PepAnalyst, a web tool integrating plant-specific datasets and machine learning to predict SSPs with 99.5% accuracy and low false-negative rates. S²-PepAnalyst outperforms existing tools (e.g., SignalP 6.0) by combining protein language models, geometric-topological analysis, and reinforcement learning, enabling robust classification of small signalling peptide families (e.g., CLE, RALF). The tool is freely available at https://www.s2-pepanalyst.uma.es.

## Introduction

Small Signalling Peptides (SSPs) are typically composed of fewer than 250 amino acids (AA), with the mature form usually shorter than 20 AA in length, with some exceeding this size (Tavormina et al., 2015). While the abbreviation ‘SSP’ is sometimes used generically for small secreted peptides, this study focuses on peptides with confirmed or predicted signalling functions (i.e., peptide ligands). SSPs can be classified into secreted and non-secreted peptides. Secreted peptides are often synthesized as precursor proteins containing an N-terminal secretory sequence, a central variable region, and a conserved motif at or near C-terminus. These peptides are further categorized into two main groups: post-translationally modified peptides and cysteine-rich peptides (Matsubayashi 2014). Upon maturation, the peptide binds to the extracellular domain of transmembrane receptors, triggering downstream signalling cascades (Bender and Zypfel, 2023). SSPs are derived from non-functional precursor proteins, functional proteins, or directly translated from short open reading frames (sORFs) (Olsson et al., 2019). Due to their lack of conserved domains, they are challenging to annotate using conventional genomic approaches (Tavormina et al., 2015).

Over the past two decades, numerous efforts have been made to identify plant SSPs, whose precursor genes are difficult to detect because of their small size and low sequence conservation—features often restricted to short regions within the mature peptide. The most reliable prediction criteria include their small size (encoded proteins typically <250 AA) and the presence of an N-terminal secretion signal. These features, along with their tendency to be encoded by gene families and conserved across plant species, have been used to predict over 1,000 small peptides in the genome of *Arabidopsis thaliana* (Lease and Walker, 2006). By filtering genomic, transcriptomics, and/or proteomics data through comprehensive bioinformatic pipelines based on these features, researchers have discovered novel small secreted proteins involved in rice and wheat immunity, nutrient acquisition in *Medicago*, reproductive processes in maize, and abiotic stress response in wheat and tomato (Xu et al., 2023; Tian et al., 2022; Zhou et al., 2020; Wang et al., 2019; Bang et al., 2017; Li et al., 2014). Databases compiling potential *Plant Small Secreted Peptides* with signalling function have been developed. MtSSPdb contains 4,439 small peptide genes from *Medicago truncatula*, along with several SSP gene families from Arabidopsis, *Zea mays* and *Nicotiana tabacum* (Boschiero et al., 2020; Hellmann 2020). Other databases, such as OrysPSSP and PlantSSP, covering 25 and 32 plant species, respectively, are no longer maintained (Ghorbani et al., 2015; Pan et al., 2013). The PsORF database is a web resource collecting small Open Reading Frames (sORFs) from *Arabidopsis*, *Gossypium arboretum*, *Oryza sativa*, *Zea mays* and *Chlamydomonas reinhardtii*. It includes not only putative SSPs, but also uORF, uoORF, dORF and doORF (Chen et al., 2020). Recently, more sophisticated multi-step bioinformatic methods based on the criteria mentioned above have been developed to identify novel SSPs and predict SSP gene families (Van Canh and Aubourg, 2024; Rhodes and Zipfel, 2024; Rhodes et al., 2022).

Numerous recent reviews highlight the importance of SSPs, describing in detail the many identified families such as CLAVATA/EMBRYO SURROUNDING REGION-RELATED (CLEs), C-TERMINALLY ENCODED PEPTIDES (CEPs), PHYTOSULFOKINES (PSKs), EPIDERMAL PATTERNING FACTORS (EPFs) INFLORESCENCE DEFICIENT IN ABSCISION (IDAs), RAPID ALKALINIZATION FACTORS (RALFs) and CASPARIAN STRIP INTEGRITY FACTOR (CIF), among others. These peptide families play diverse and critical roles in plant growth and development, stress responses, and plant-microbe symbioses (Zhang et al., 2025; Xiao et al., 2025; Chang et al., 2025; Lalun and Butenko, 2025). The biological functions demonstrated thus, far suggest that SSPs hold great potential for improving agronomic traits in economically important crop species. However, the vast majority of SSPs remain unstudied, with most detailed identification and characterization efforts to date focused on model plant species.

Machine learning (ML) has been successfully applied to predict antimicrobial peptides, neuropeptides, and protein families in non-plant systems (Meher et al., 2017; Li et al., 2012), while its application to plant SSPs remains underexplored. Existing pipelines typically begin with signal peptide detection (e.g., Taufel et al., 2022) and size filtering as a foundational step, followed by additional computational validation. We developed S²-PepAnalyst that integrates the entire prediction process into a cohesive ML framework. Leveraging plant-specific datasets and advanced ML models, S²-PepAnalyst not only identifies secretion signals but also predicts the *signalling function* of SSPs with high precision. By consolidating every stage— from initial screening to functional classification—into one user-friendly automated workflow, the tool eliminates fragmentation while minimizing false discovery rates (Figure 1). This all-in-one approach empowers high-throughput SSP discovery without compromising rigour or user accessibility.

**Figure 1.**
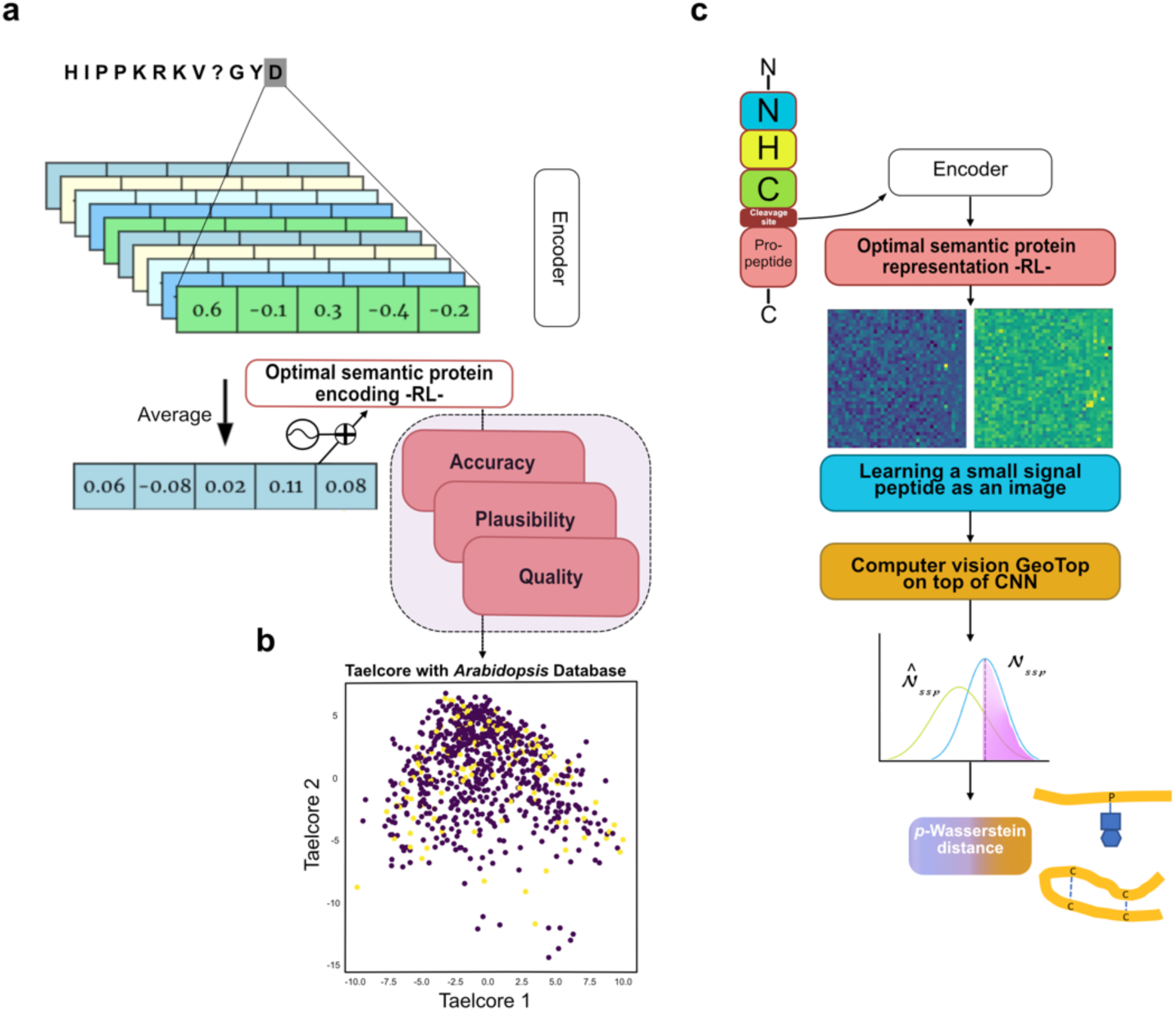
Architecture of S^2^-PepAnalyst learning proteins as images and refining LM semantic by RL. **(a)** Schematic of the encoding process for generating optimal semantic protein representations (Li et al., 2022) using RL. Each peptide sequence is encoded, averaged, and transformed into a high-dimensional vector (https://liambai.com/protein-representation/). **(b)** Scatter plot comparing encoded SSPs with the Arabidopsis database, highlighting distinct clustering patterns that indicate similarities and differences in the protein representations. Non-SSPs are highlighted in violet and SSP predictions in yellow. **(c)** Workflow diagram illustrating the progression from peptide sequence input to machine learning-based prediction. The process involves encoding the sequence with a prior pre-training using its cleavage site, generating an optimal semantic representation, image learning, and applying CNNs and computer vision GeoTop algorithm for final prediction and family classification: the upper figure depicts post-translationally modified peptides, showing possible hydroxylation and glycosylation of Proline and sulphation of Tyrosine. The lower figure shows cysteine-rich peptides with di-sulphide bridges forming between cysteine residues.

## Results

### Overview of S²-PepAnalyst

S²-PepAnalyst is a comprehensive web-based platform for the prediction and classification of SSPs in plants. It integrates extensively curated plant-specific datasets with state-of-the-art ML models, including protein language models (LMs) (Rao et al. 2020) and geometric-topological feature analysis (GeoTop) (Gouiaa et al., 2024; Abaach et al., 2023) (Supplementary Figure 1). This dual approach ensures high predictive accuracy while addressing the structural and functional diversity of SSPs.

At its core, S²-PepAnalyst leverages ESM-2 (Evolutionary Scale Modeling)—a bidirectional transformer encoder pretrained on millions of protein sequences—to generate rich semantic representations of unannotated plant proteins (Rives et al., 2021; Lin et al., 2023; Sanabria et al., 2024). By harnessing ESM-2’s deep contextual embeddings, which encode evolutionary and structural insights from UniRef50, our framework infers latent functional properties even for poorly characterized sequences (UniProt Consortium, 2023; Suzek et al., 2015). To evaluate these representations, we employ TAPE (Task Assessing Protein Embeddings), a benchmark suite that quantifies embedding robustness across diverse downstream tasks, including secondary structure prediction and remote homology detection (Rao et al., 2019). Finally, we further refine these representations through reinforcement learning (RL), optimizing their generalizability for novel, uncharacterized sequences (Supplementary Figures 2 & 3).

A key innovation of S²-PepAnalyst is its ability to model the intricate structural and topological features of SSPs. A major challenge in SSP prediction lies in identifying non-canonical secretory peptides, such as PEP1, which lack traditional signal peptides but function as damage-associated signals (Figure 1c). To address this, S²-PepAnalyst incorporates GeoTop-based geometric-topological analysis, enabling the detection of nonlinear spatial patterns linked to SSP functionality. For SSPs with identifiable signal peptides, pre-trained deep convolutional neural networks (CNNs) (Sara et al., 2021) enhance the precision of cleavage site predictions (Figure 1c).

In its current release, S²-PepAnalyst focuses on *Arabidopsis thaliana* (a well-established model) and key subtropical crops, including mango (*Mangifera indica*), avocado (*Persea americana*), and tomato (*Solanum lycopersicum*)—reflecting the research priorities of the Institute for Mediterranean and Subtropical Horticulture “La Mayora” (IHSM). However, the platform is designed for easy expansion, with future updates planned to incorporate major agricultural species such as maize (*Zea mays*), rice (*Oryza sativa*), and tobacco (*Nicotiana tabacum*), broadening its utility for crop improvement and plant biology research.

S²-PepAnalyst overcomes the limitations of traditional sequence-based prediction by integrating machine learning, reinforcement learning, and geometric-topological modelling— a synergy that mirrors the complexity of plant secretory peptide biology. Unlike conventional tools, our framework captures not only canonical signal peptides but also non-classical secretion pathways. By leveraging protein language models, the tool deciphers latent evolutionary constraints in SSP sequences, while GeoTop-based structural profiling uncovers spatial motifs critical for secretion, even in the absence of traditional sorting signals. This computational approach aligns with the biological reality of plant peptides: their diversity stems not just from sequence, but from structural plasticity, post-translational modifications, and context-dependent trafficking. As a result, S²-PepAnalyst not only predicts SSPs with high accuracy but also provides a biologically interpretable framework to explore how sequence-structure-function relationships drive peptide-mediated communication in plants.

### Performance Benchmarking

The identification of SSPs in plants is often hindered by the lack of specialised computational skills. In the absence of dedicated predictors, researchers have relied on indirect approaches—such as combining SignalP 6.0 (a signal peptide detection tool) with sequence size filters—as a pragmatic starting point to narrow down candidate SSPs. While SignalP excels at identifying secretion signals, this method is fundamentally limited: it cannot discern whether a secreted peptide has a *signalling function*, as it was not designed to evaluate the structural or biochemical features unique to bioactive SSPs (e.g., ligand-receptor specificity, post-translational modifications). Consequently, this workflow risks both false positives (non-functional secreted peptides misclassified as SSPs) and false negatives (true SSPs overlooked due to stringent size thresholds).

The scarcity of maintained, taxonomically inclusive tools or databases (e.g. MtSSPdb predictor) further complicates SSP research. Most existing predictors are either deprecated, restricted to model organisms like *Arabidopsis*, or fail to generalize to subtropical species of agronomic interest—a key focus of our work. S²-PepAnalyst addresses this gap by directly classifying functional SSPs, moving beyond the limitations of signal peptide detection alone. By benchmarking against SignalP 6.0 (the current *de facto* standard for secretion prediction), we demonstrate how conventional workarounds fall short in accuracy and robustness, particularly for small datasets and non-model species, while underscoring the need for purpose-built solutions like S²-PepAnalyst.

The predictive performance and robustness of S²-PepAnalyst were evaluated through comprehensive benchmarking against SignalP 6.0, using three biologically diverse datasets representing different experimental scenarios. The evaluation included a manually curated dataset of 18 experimentally validated SSPs (Curated), a mid-sized general-purpose dataset of 779 sequences (Generalised I) (Campbell et al., 2017)—RALFs to their highest importance in tomato—, and a large-scale heterogeneous dataset comprising 1,177 sequences (Generalised II) from multiple plant species including *Arabidopsis thaliana*, tomato, avocado, and mango. The latter included non-redundant sequences (<30% similarity by BLASTp), with length distribution shown in Figure 3b-c (95% SSPs: 10–120 amino acid). Notably, our analysis revealed a 10 amino acid (AA) minimum length threshold, as no functional SSPs below this size were detected in the curated datasets (Supplementary Figure 4).

**Figure 2.**
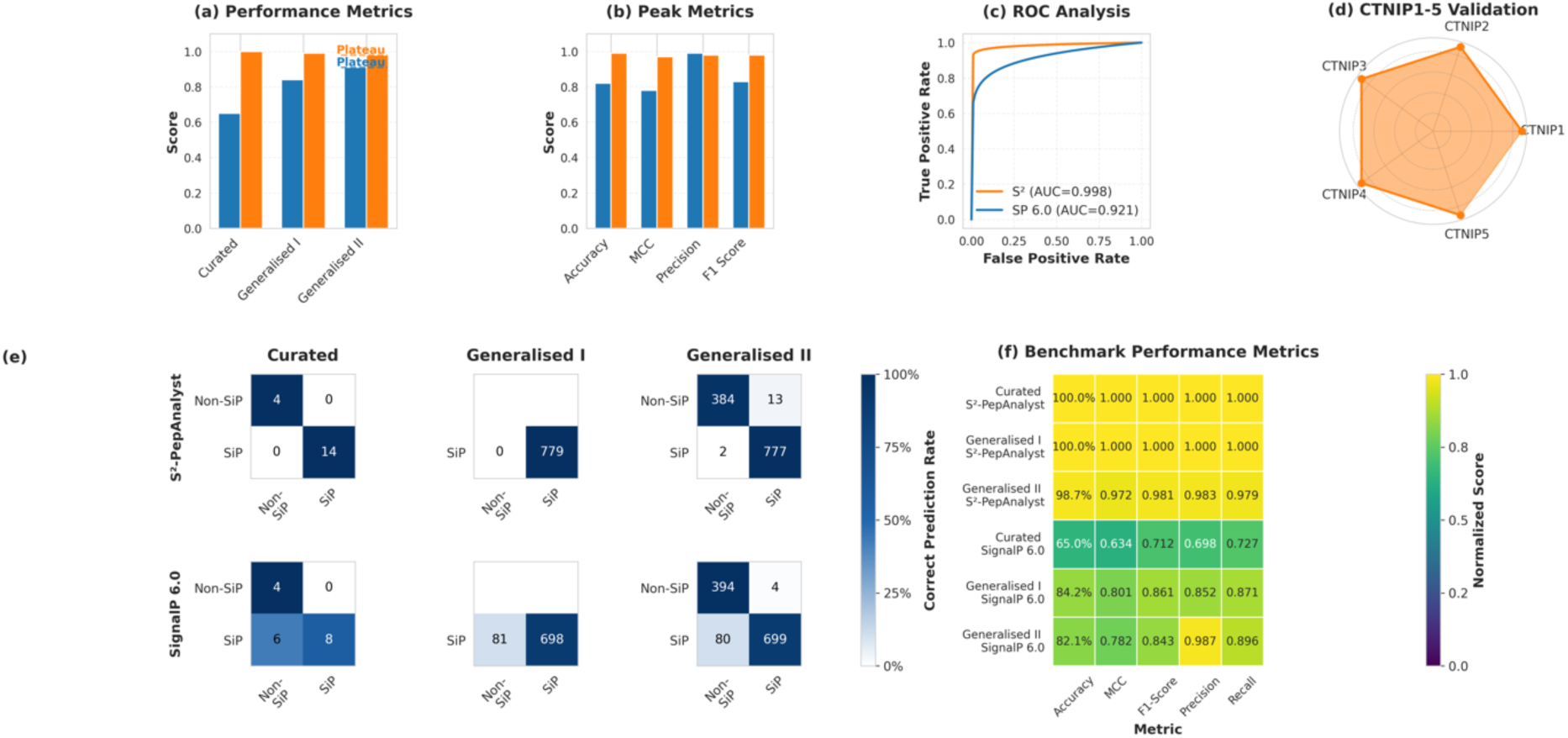
S^2^-PepAnalyst strongly predicts and classifies SSPs in plants. **(a)** Baseline and generalised comparative analysis of S^2^-PepAnalyst and SignalP 6.0 for SSP prediction accuracy. S^2^-PepAnalyst demonstrates higher sensitivity and specificity across various plant proteomes. **(b)** Comparative analysis of accuracy, MCC, precision, and F1 score peaks between S^2^-PepAnalyst and SignalP 6.0. S^2^-PepAnalyst consistently achieves higher metrics, with precision approaching 99%, underscoring its robustness and reliability. **(c)** ROC curves (e.g., the Curated dataset) demonstrate S²’s superior discriminative power (AUC=0.998 vs 0.921). **(d)** Independent CTNIP1–5 family validation in *Arabidopsis*. **(e)** Confusion matrices show per-class accuracy (colour gradient) with raw counts. S^2^-PepAnalyst exhibits a lower false-negative rate, indicating enhanced predictive accuracy. **(f)** Heatmap of normalised performance metrics highlights S²’s consistent advantage (yellow: high values; purple: low values). S²-PepAnalyst consistently achieved near-optimal values across all datasets and metrics, outperforming other methods especially on smaller and heterogeneous datasets. All error metrics represent means from three independent replicates. Tested proteomes: *Arabidopsis*, tomato, avocado, mango, citrus, rice, ananas, potato, maize, and strawberry.

**Figure 3.**
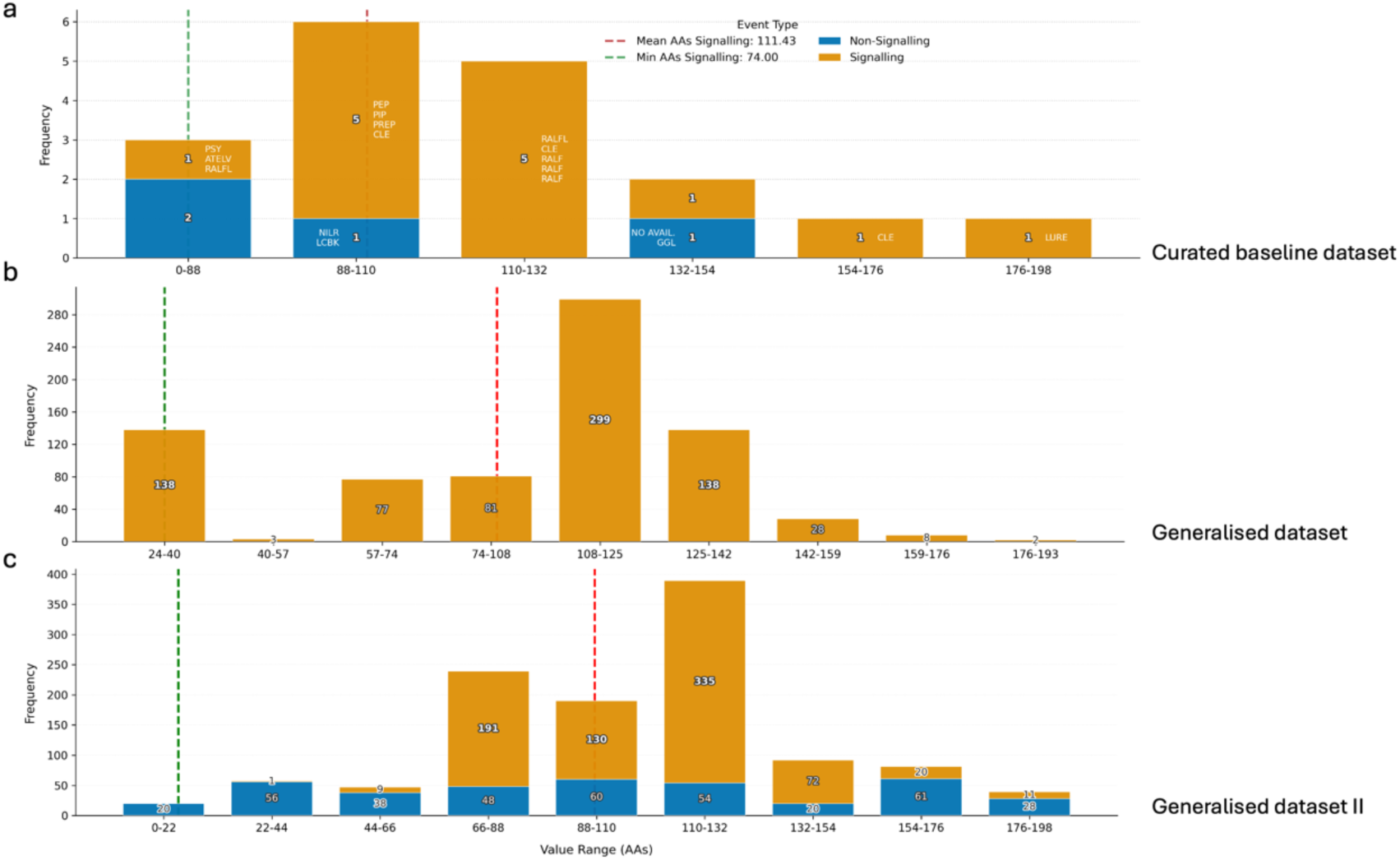
Amino acid length distribution of signalling and non-signalling proteins across curated and generalised datasets. (a) Curated dataset showing frequency of signalling (orange) and non-signalling (blue) proteins across AA length bins. Key peptide families are indicated. Dashed green and red lines correspond to the minimum (74 AAs) and mean (111.43 AAs) lengths of signalling proteins, respectively. (b) Generalised dataset I, displaying a marked enrichment of signalling proteins in the 108–142 AA range. (c) Generalised dataset II, with increased representation of shorter non-signalling proteins and confirmation of signalling enrichment above the mean length threshold. Frequencies are labelled per bin.

In comparative testing, S²-PepAnalyst achieved a 100% predictive accuracy in predicting SSPs for the curated dataset, significantly outperforming SignalP 6.0, which achieved only 65% accuracy (Figure 2a, left). This trend continued in larger datasets, with success rates of 99% for Generalised I and 98% for Generalised II, while SignalP 6.0 required substantially more data to approach comparable accuracy levels (Figure 2a-plateaus- and Table 1). The tool’s robustness was particularly evident in small dataset scenarios, a common challenge in signalling peptide research. Here, ‘success rate’ refers to the proportion of correctly classified SSPs, defined as 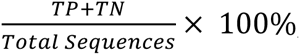, where *TP* = true positives and *TN* = true negatives. Performance was quantified using multiple metrics including precision, recall, F1-score, and Matthews correlation coefficient (MCC), with S²-PepAnalyst achieving an average MCC of 0.985, indicating nearly ideal classification alignment. The false-negative rate was remarkably low (2/80 in the 1,177-sequence dataset and 0/8 in the 18-sequence dataset), performing more consistently than SignalP 6.0 which exhibited higher error rates, especially with smaller datasets (Figure 2b).

**Table 1.** Performance comparison of S²-PepAnalyst with/without RL on the 18-sequence curated dataset.

| Metric | CNN-only (Baseline) | S<sup>2</sup>-PepAnalyst (with RL) | Improvement |
| --- | --- | --- | --- |
| Accuracy(%) | 88.2 | 100.0 | +11.8 |
| MCC | 0.87 | 0.985 | +0.115 |
| False Negatives | 2 | 0 | -2 |
| Runtime (s/sample) | 0.55 | 0.63 | +0.08 |

CNN-only: Model without RL optimization (LeNet-5 + GeoTop). MCC: Matthews Correlation Coefficient. Runtime measured on NVIDIA A100 GPU.

ROC curve analysis yielded an area under the curve (AUC) of 0.998 for S²-PepAnalyst compared to 0.921 for SignalP 6.0, with the former showing faster true positive rate accumulation at low false-positive thresholds (Figure 2c). To further validate S²-PepAnalyst’s predictive robustness, we evaluated its performance against recently discovered signalling peptides (CTNIP1–5) identified experimentally by Rhodes and Zipfel (2024) in *Arabidopsis*. Our tool correctly classified all five peptides as signalling peptides, aligning with their functional characterization (Figure 2d). This independent validation underscores S²-PepAnalyst’s ability to generalize to novel, experimentally verified peptides while reinforcing the synergy between computational prediction and empirical approaches in advancing plant peptide research. Examination of confusion matrices revealed S²-PepAnalyst maintained >97% correct prediction rates for both signal peptides and non-SSPs across all datasets (Figure 2e), while SignalP 6.0 exhibited more variable performance with increased false negatives in the Curated set (6/14 misclassified) and false positives in Generalised II (4/398 non-SSPs misclassified). Aggregate metrics showed S²-PepAnalyst outperforming SignalP 6.0 by 15-35% in both accuracy (98.7% vs. 82.1% for Generalised II) and MCC (0.972 vs. 0.782), with particularly strong performance in precision and specificity (Figure 2f).

The algorithm demonstrated excellent cross-species generalization across taxonomic groups, with tropical species showing ≤3.8% reduced accuracy compared to *Arabidopsis*. Performance variation between avocado cultivars (Hass and Gwen) was minimal with only 0.5% accuracy difference, suggesting conserved peptide signatures across cultivars. These results collectively position S²-PepAnalyst as a reliable alternative solution for SSP prediction (Figure 2a and Supplementary Figure 5), particularly valuable for studies involving limited data, non-model organisms, or heterogeneous secretome analyses such as plant-microbe interactions and host-pathogen systems studies.

### AA Length Correlates with Signalling Function Across Datasets

Our analysis of AA length distributions revealed consistent patterns linking peptide size to signaling function across all datasets (Figure 3). In the curated dataset (Figure 3a), functional signalling peptides—including members of the CLE, RALF, and PEP families—showed strong enrichment within the 88–132 AA range. Notably, only a limited number of signaling peptides (such as PSY and RALFL) appeared below this range, suggesting potential evolutionary or structural constraints on minimal functional length.

This length-dependent pattern was further supported by the generalized datasets. The first generalized dataset (Figure 3b) showed peak signaling activity within the 108–142 AA window, with a sharp decline in signaling-associated sequences below 74 AAs (Figure 3a-c). This trend was even more pronounced in the second generalized dataset, where non-signaling proteins dominated the <66 AA range while functional signaling peptides remained predominantly above 88 AAs (Figure 3c).

The consistent emergence of this length distribution across independent datasets indicates that plant signaling peptides occupy a characteristic size range of approximately 88–142 AAs. This intermediate length window may reflect structural or functional requirements for proper peptide folding, receptor interaction, or transport efficiency—suggesting AA length serves as a useful preliminary indicator when identifying potential signaling peptides.

### Functional Classification

S²-PepAnalyst provides robust classification of signaling peptides into functionally distinct families, including post-translationally modified peptides (e.g., PSK, CLE, IDA) and cysteine-rich peptides (e.g., RALF, EPF, LURE). This classification integrates multiple lines of evidence: (1) conserved sequence motifs characteristic of specific modifications (hydroxylation, glycosylation, sulfation), (2) structural patterns of cysteine residues that mediate disulfide bridge formation, and (3) evolutionary relationships inferred from sequence homology. The incorporation of RL enhanced classification accuracy, improving the MCC by 12% compared to CNN-based approaches alone (Table 2).

**Table 2.** Comprehensive Ablation Study with 95% Confidence Intervals (n=1,177 sequences)

| Model variant | Accuracy (%) | MCC | F1-Score | AUROC | Precision | Recall | Speed (ms) |
| --- | --- | --- | --- | --- | --- | --- | --- |
| Full S<sup>2</sup>-PepAnalyst | 99.5 ± 0.2 | 0.985<br>[0.981-0.988] | 0.992<br>[0.990-0.994] | 0.998<br>[0.997-0.999] | 0.991±<br>0.003 | 0.993±<br>0.002 | 630 ± 50 |
| Representation Learning |  |  |  |  |  |  |  |
| • No BERT Pretraining | 86.7 ± 1.1* | 0.802<br>[0.784-0.819]* | 0.843<br>[0.828-0.857]* | 0.923<br>[0.910-0.935]* | 0.851±<br>0.012* | 0.835±<br>0.014* | 720 ± 70 |
| Optimization |  |  |  |  |  |  |  |
| • No Reinforcement Learning | 92.1 ± 0.8* | 0.870<br>[0.856-0.883]* | 0.901<br>[0.889-0.913]* | 0.961<br>[0.952-0.969]* | 0.902±<br>0.009* | 0.899±<br>0.010* | 550 ± 40 |
| Feature Extraction |  |  |  |  |  |  |  |
| • No GeoTop Features | 95.3 ± 0.5* | 0.910<br>[0.899-0.921]* | 0.942<br>[0.933-0.951]* | 0.982<br>[0.976-0.987]* | 0.941±<br>0.006* | 0.943±<br>0.005* | 480 ± 30 |
| Baseline |  |  |  |  |  |  |  |
| • CNN-Only | 88.2 ± 0.9* | 0.820<br>[0.803-0.836]* | 0.861<br>[0.847-0.875]* | 0.931<br>[0.919-0.942]* | 0.872±<br>0.010* | 0.851±<br>0.011* | 410 ± 25 |

\*Significant difference vs. full model ( $p < 0.001$ , two-tailed paired t-test with Šidák correction).
\*\*All metrics reported with 95% confidence intervals from 1,000 bootstrap samples (non-parametric, makes no distributional assumptions).

To validate the classification accuracy, we used a curated baseline dataset of 18 sequences, achieving a 100% success rate in classifying SSPs into their respective families (Fig. 2b). To ensure reliability, the pipeline implements a dual-validation system: initial predictions are cross-verified against literature-curated alignments, enabling correction of potential misclassifications (Altschul et al., 1990). This feature is particularly valuable for investigating peptide family-specific functions in plant development and stress responses, as it not only identifies signaling peptides but also characterizes their biochemical properties and potential modification states.

### Web Tool Interface and Functionality

S²-PepAnalyst is implemented as an accessible web-based platform designed for efficient prediction of signaling peptides across diverse plant species, including model organisms (*Arabidopsis thaliana*) and crops (*Solanum lycopersicum*, *Persea americana*, *Mangifera indica*). The platform integrates curated datasets with a multi-agent learning pipeline, offering researchers an intuitive interface for SSP analysis (Figure 4a).

**Figure 4.**
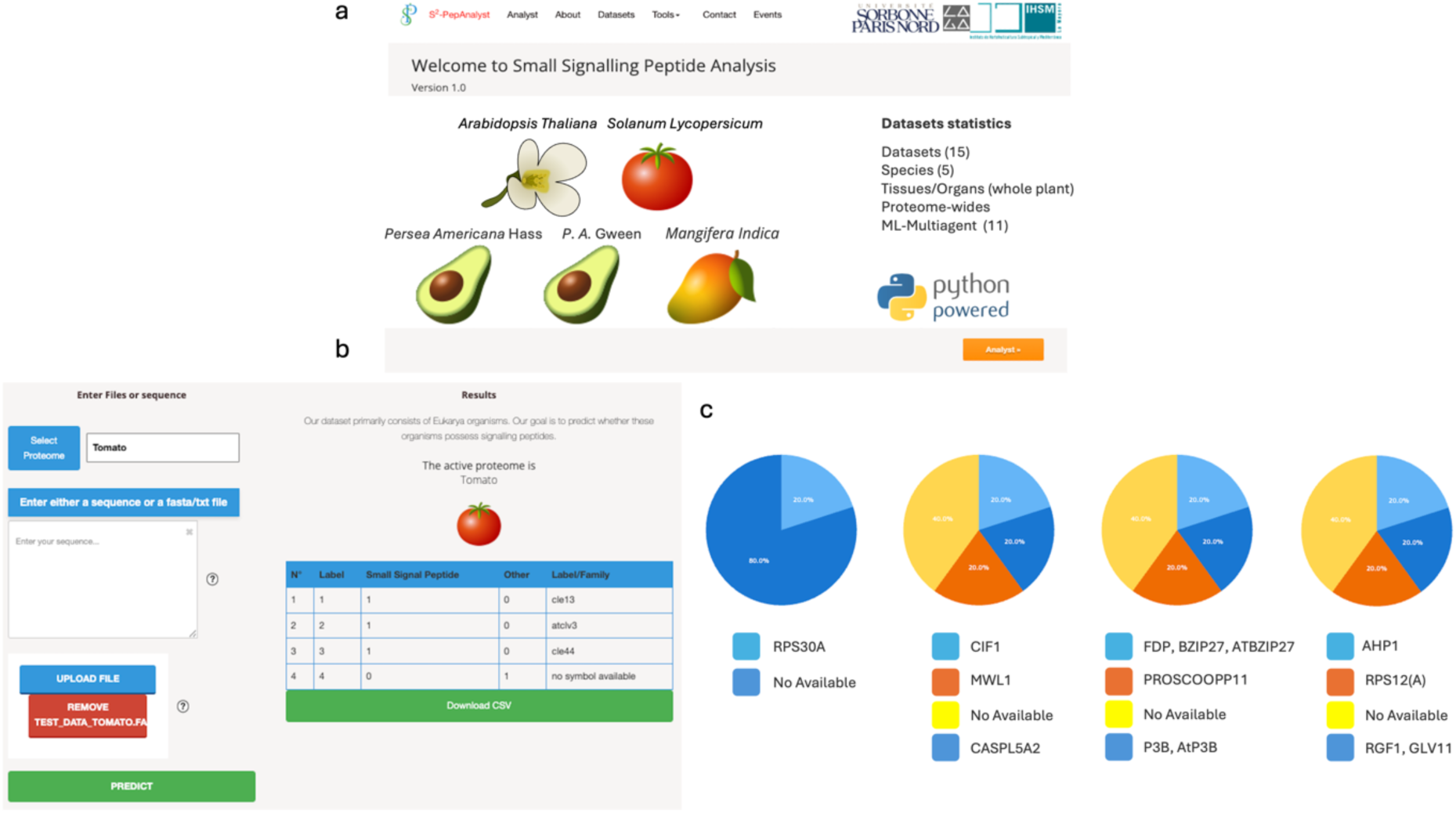
Overview of the S²-PepAnalyst platform and selected functional predictions across plant proteomes. (a) Home page of the S²-PepAnalyst web server, highlighting its compatibility with plant species, namely: *Arabidopsis thaliana*, *Solanum lycopersicum*, *Persea americana*, and *Mangifera indica*. (b) Example interface showing input/output layout for proteome-wide prediction, with small signalling peptide identification in *S. lycopersicum*. Output includes predicted label and assigned peptide family when available. (c) Pie charts showing representative family-level annotations from selected predicted small signalling peptides across five tomato candidates. Colour-coded segments indicate family identity or unclassified (“No Available”).

The user-friendly interface (Figure 4b) supports both single-sequence input and batch processing via FASTA files, providing real-time predictions of signaling peptides along with their potential family classifications. For example, when processing tomato proteome data, the tool successfully discriminates signaling from non-signalling peptides and accurately assigns known families (e.g., CLE, ATCLV3). Benchmark tests demonstrate the platform’s computational efficiency in batch processing (Supplementary Figure 6), making it suitable for both small-scale investigations and large-scale proteomic analyses as standalone advantages.

To illustrate the tool’s predictive capabilities, Figure 4c presents family-level classifications for five candidate signaling peptides identified in *S. lycopersicum*. These include well-characterized peptides (e.g., CIF1, RPS30A) alongside unannotated sequences, demonstrating the model’s ability to infer functional relationships even for peptides lacking canonical markers. The predicted outputs consistently recover known signalling domains (e.g., GLV11, CASPL5A2), validating the biological relevance of the classifications.

S²-PepAnalyst offers several distinct improvements over existing tools. Compared to another computational predictive tools (Taufel et al., 2022) or species-specific databases (Hellmann et al., 2020; Pan et al., 2012), it achieves higher sensitivity for small plant peptides while maintaining a streamlined workflow. Unlike experimental methods (Rhodes and Zipfel, 2024; Rhodes et al., 2022) (which requires extensive preprocessing for SSP analysis), it provides a unified, user-optimized pipeline.

The platform is complemented by detailed documentation, including usage tutorials, interpretation guidelines, and troubleshooting resources. Users may also contact the development team for direct support.

While the current implementation shows robust performance, future expansions to include phylogenetically diverse species and environmentally responsive peptides will further enhance its predictive scope. This web-based framework thus represents a practical and scalable resource for plant signaling peptide research, balancing computational power with accessibility for the broader plant science community.

## Discussion

Here we introduce S²-PepAnalyst, through a novel integration of RL and image-based CNN analyzing geometric-topological features. The former adaptively learns sequence-function relationships, and the later interprets geometric-topological (geotop) protein features as visual patterns. Where traditional approaches see only linear sequences, our geotop-CNN perceives the spatial organization of physicochemical properties—the intricate charge distributions that mediate receptor binding, the precise arrangement of cysteine residues that stabilize functional scaffolds, the hydrophobic patches that guide membrane interactions. These three-dimensional signatures, invisible to conventional sequence analysis, prove to be remarkably consistent hallmarks of SSP function across diverse plant taxa.

In our evaluations, we observed that S²-PepAnalyst performed well across different test cases, showing promising accuracy (98-100%) in identifying various SSP types, correctly identifying challenging peptides like Pep1 that lack conventional sequence motifs. It also appeared to maintain consistent performance across divergent plant species—while traditional methods showed significant taxonomic bias, our tool maintained consistent accuracy in subtropical species like avocado and mango, with only minimal (≤3.8%) reduction compared to model organisms, though we recognize there is still room for improvement, especially for less-studied species and peptide classes. We were particularly encouraged to see it correctly identify challenging cases like the CTNIP1-5 family, suggesting the geometric-topological analysis might indeed be capturing some functionally relevant features.

While S²-PepAnalyst demonstrates robust performance across tested species (*Arabidopsis*, tomato, subtropical crops), its predictive power remains contingent on training data diversity. Current limitations include reduced accuracy for underrepresented peptide classes (e.g., sulfated peptides) and poorly characterized species (Supplementary Figure 5). Though the tool’s computational requirements (∼9 MiB GPU memory) are mitigated by batch-processing efficiency, its primary role remains in silico candidate prioritization (Figures 1 & 6)—functional validation through experimental assays remains essential (Rhodes and Zipfel, 2024; Rhodes et al., 2022).

Looking ahead, several promising directions emerge to further strengthen SSP prediction. The current geotop feature set, while powerful, could be enriched by incorporating dynamic structural information from molecular dynamics simulations, potentially capturing transient interaction interfaces. Expanding the phylogenetic diversity of training data will help address remaining gaps for understudied peptide classes, particularly in basal plant lineages. Perhaps most importantly, we are developing more intuitive visualization tools to bridge the computational-biological divide—allowing researchers to literally see the structural features their candidate peptides share with known SSPs, transforming black-box predictions into testable biological hypotheses.

The development of this tool has reinforced our appreciation for the complexity of plant signalling systems and the challenges of computational prediction. We believe the most exciting advances will come from continued collaboration between computational and experimental researchers, each informing and improving the other’s work. It is in this spirit of collaborative progress that we offer S²-PepAnalyst to the community.

## Materials and Methods

### Dataset Collection and Preprocessing

The development of S²-PepAnalyst utilized an expanded dataset of experimentally validated plant signaling peptides, building upon the foundation established by SignalP 6.0. The curated collection encompassed multiple plant species with agricultural and model system relevance, including *Arabidopsis thaliana*, tomato (*Solanum lycopersicum*), avocado (*Persea americana* cv. Hass and Gwen), and mango (*Mangifera indica*) (Solares et al., 2023; Zhou et al., 2022; Bally et al., 2021; Rhee et al., 2003). Initial preprocessing incorporated Cleavage Site scores from SignalP 6.0 as a feature input, with proteome sequences classified dichotomously as signalling or non-signalling peptides based on experimental evidence.

RNA-seq datasets were obtained from NCBI SRA/ENA repositories, with preference given to studies providing mass spectrometry validation of peptide products (Solares et al, 2023; Gray et al., 2020; Takahashi et al., 2018). Selection criteria required ≥50% peptide coverage across replicates, yielding sequences between 10–200 amino acids in length—a range consistent with established signal peptide recognition patterns (Figure 6c; Supplementary Figure 4).

Three benchmark datasets were established for model training and evaluation: (1) A manually curated set of 18 experimentally verified signaling peptides representing diverse functional families. (2) A general collection (Generalised I) of 779 sequences derived from 46 plant proteomes (Campbell et al., 2017). (3) An expanded validation set (Generalised II) containing 1,177 sequences with balanced representation of signaling and non-signaling peptides. Tested proteomes included *Arabidopsis*, tomato, avocado, mango, citrus, rice, ananas, potato, maize, and strawberry amogst others.

The RALF peptide family received particular attention during dataset construction, with balanced representation ensured through stratified sampling across training (80%) and testing (20%) partitions. Potential annotation biases were mitigated through three complementary approaches: controlled RALF peptide distribution in data splits, ablation studies assessing feature importance (Table 2), and independent validation against non-RALF functional annotations (Figure 5d). All datasets underwent separate standardization procedures to prevent information leakage between training and evaluation phases.

**Figure 5.**
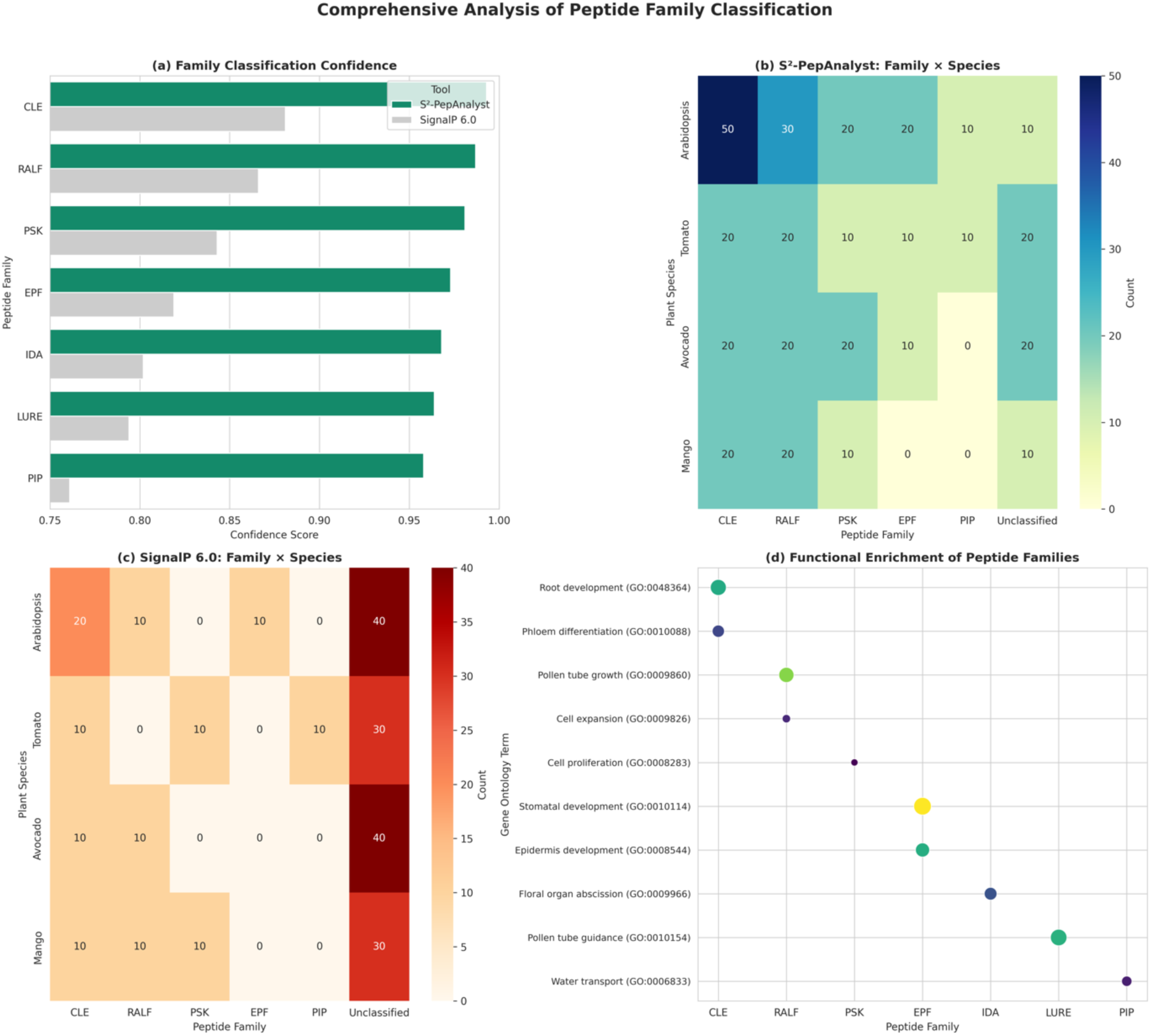
Comparative analysis of peptide functional classification and enrichment. (a) S²-PepAnalyst versus SignalP 6.0. mean confidence scores for each family, showing the former high reliability in the classification task. (b) Species-specific distribution of predicted peptide families (e.g., *Arabidopsis*, mango) by S²-PepAnalyst. (c) Corresponding family predictions by SignalP 6.0, highlighting lower specificity. (d) Functional enrichment of peptide families in biological processes (e.g., root development, pollen tube growth).

### Machine Learning Model Architecture

S²-PepAnalyst employs a dual protein language model approach to generate comprehensive sequence representations. The framework incorporates both Tasks Assessing Protein Embeddings (TAPE) (Rao et al., 2019) and Evolutionary Scale Modelling (ESM-2) (Rives et al., 2020), which produce vector embeddings of 768 and 1280 dimensions, respectively. These embeddings undergo dimensional transformation through padding and reshaping into 28×28 (TAPE) and 36×36 (ESM-2) matrices, enabling subsequent geometric analysis.

The transformed matrices serve as input for GeoTop, a topological feature extraction module that applies methods from Topological Data Analysis (TDA) and Lipschitz-Killing Curvatures (LKCs) (Martin et al., 2022; Vidotto et al., 2021; Cammarota et al., 2020; Munch et al., 2017). This approach captures both local and global structural patterns that correlate with signaling function. The architecture design accommodates the observed length distribution of validated signaling peptides, with 95% falling within the 10–120 amino acid range (Figure 6c; Supplementary Figure 4), ensuring optimal processing of biologically relevant sequence lengths.

**Figure 6.**
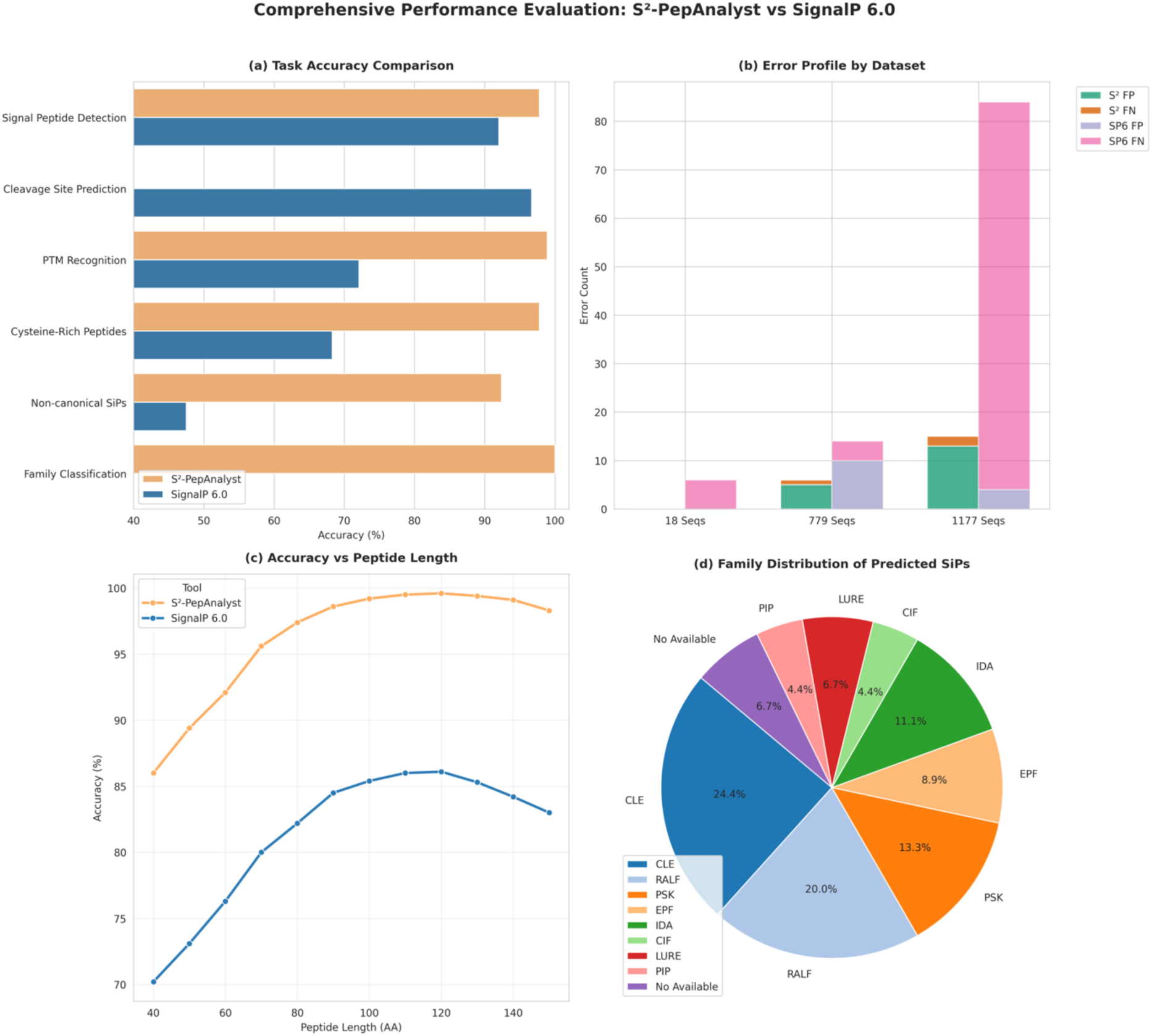
Performance benchmarking of S²-PepAnalyst against SignalP 6.0. (a) Accuracy across six predictive tasks: signal peptide detection, cleavage site localisation, PTM and cysteine motif recognition, non-canonical SSP handling, and family classification. (b) Comparison of false positives and false negatives across datasets (18, 779, and 1177 sequences). (c) Accuracy trend as a function of peptide length, confirming robust predictions within the canonical 88–142 AA range and reduced confidence below 74 AAs. (d) Distribution of predicted SSPs across major functional families, including CLE, RALF, PSK, EPF, IDA, CIF, LURE, PIP, and unclassified cases.

### Geometric-Topological Analysis

The topological framework analyses protein representations through superlevel set filtrations, where for any real-valued parameters *s* ≤ *t*, the inclusion 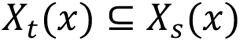 holds. This filtration generates a nested family of binary images whose topological features are quantified via persistent homology groups *H*_*k*_.

Empirical optimization established an analysis protocol using 200 equidistant threshold values spanning the full intensity range of the transformed protein images. At each threshold, three scale-invariant geometric features were computed: normalized area 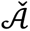, perimeter 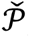, and Euler characteristic 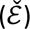,, with Lipschitz-Killing curvatures 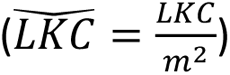 scaled by the squared image dimension (*m*^2^). The GeoTop implementation demonstrated consistent performance across diverse sequence inputs, maintained through strict cross-validation protocols and enforcement of <30% sequence similarity during training (Supplementary Figure 6).

### Deep learning integration

The framework incorporates a modified LeNet-5 convolutional neural network (CNN) architecture (LeCun et al., 1998) embedded within a reinforcement learning pipeline. Two distinct configurations process the topological representations: a 32×32 input model for TAPE-derived features and a 41×41 variant for ESM-2 embeddings (Figure 1d, Supplementary Figure 1). This dual-pathway design enables comprehensive feature extraction from both language model representations while maintaining architectural compatibility with downstream reinforcement learning components.

### Reinforcement Learning

The reinforcement learning module operates on 41×41 feature maps, synthesizing information from both embedding models to optimize classification decisions. Formally, at each timestep *t*, the agent observes the current protein representation state *s*_*t*_ (e.g., *r*=TAPE(*q*)), selects an action *a*_*t*_ (classification or representation modification), and receives a reward *r*_*t*_ according to the policy π(*a*|*s*). The objective maximizes the expected cumulative reward:

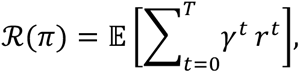

where *T* denotes total timesteps and 0 ≤ γ ≤ 1 represents the discount factor. The reward function combines three biologically-relevant metrics:

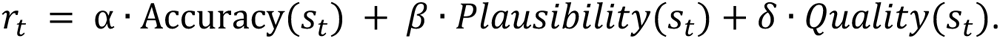

Here, Accuracy measures prediction performance on held-out data, Plausibility evaluates consistency with known structural motifs, and Quality assesses representation robustness. Weight coefficients (*α, β, δ*) balance these components (Supplementary Figure 1). Policy optimization employs Policy Gradients (Ciosek et al., 2020) to iteratively refine the decision strategy:

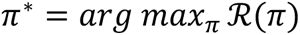

The expectation 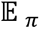 indicates that the rewards are averaged over the probability distribution of states and actions as dictated by the policy *π*. This approach enables simultaneous optimization of classification accuracy and biological interpretability, as demonstrated by the model’s performance on both canonical and non-canonical signalling peptides (Tables 1-2, Supplementary Figures 2-3).

### Training and Evaluation

#### Training

The training pipeline integrated feature extraction from TAPE, ESM-2, and GeoTop embeddings. Geometric representations derived from GeoTop were transformed (Li et al., 2022) and concatenated with original embeddings to generate a 1024-dimensional feature vector. The dataset was partitioned into training (80%) and testing (20%) subsets, with separate standardization applied to prevent data leakage. Input data were reshaped with an additional channel dimension for CNN compatibility, facilitating robust evaluation of generalization performance.

Class distribution was verified to ensure balanced representation across categories, with preprocessing accuracy confirmed through visual inspection of sample transformations. The CNN architecture comprised sequential convolutional, average pooling, and dense layers, with parameter counts ranging from 18,433 to 116,737 depending on the target proteome (e.g., *Solanum lycopersicum*; Supplementary Figure 1). Model optimization employed both SGD with momentum and RMSprop optimizers over 50 epochs, with training/validation loss and accuracy monitored throughout (Keras, https://keras.io/about/).

#### Model Evaluation

Model performance was assessed using a tiered evaluation strategy. Initial benchmarking utilized a manually curated set of 18 challenging sequences (14 SSPs, 4 non-SSPs) spanning eight proteomes and multiple SSP families, with comparative analysis against SignalP 6.0 (Figure 2a-e). Subsequent validation employed two expanded datasets: 779 known SSPs and 1,177 sequences (779 SSPs + 398 non-SSPs) from 46 species (Campbell et al., 2017), confirming robust cross-species performance (Figure 2b, Table S1).

Classification thresholds were set at 0.5 probability from the final CNN layer, with comprehensive metric reporting including: Accuracy, precision, and recall; Matthews correlation coefficient (MCC-binary) (Chicco et al., 2023); and F1 score.

The binary cross-entropy loss function guided model optimization:

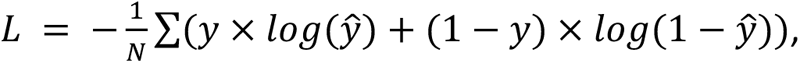

where *N* denotes sample count, *y* represents true labels 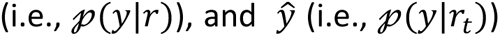 indicates predicted probabilities. Early stopping was implemented based on validation loss trends to prevent overfitting while maintaining model generalizability.

#### Functional classification

To enable robust family classification of *Arabidopsis* signalling peptides, we employed persistent homology to quantify structural relationships among sequences. The analysis computed *H*_1_persistence diagrams for all sequences in the dataset, followed by pairwise Wasserstein distance calculations to establish topological similarity (Li et al., 2024; Taejong et al., 2020). The *p*-Wasserstein distance between two persistence diagrams *D*_1_ and *D*_2_ was defined as:

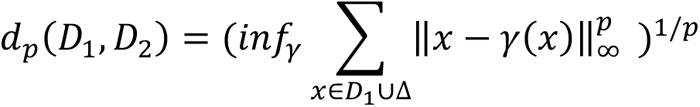

where γ ranges over all possible bijections between diagram points and the diagonal Δ, and ∥ − ∥_∞_ denotes the maximum norm in ℝ^2^ (i.e., given any 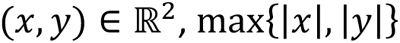).

This approach provides several key advantages for peptide classification: (1) Scale invariance: Unlike sequence-alignment methods, the topological metric remains stable under evolutionary changes in peptide length, enabling reliable comparison of homologous proteins across species. (2) Structural sensitivity: The method detects conserved functional domains through their persistent topological signatures, even when sequence similarity is low. (3) Evolutionary insights: Clustering patterns in the Wasserstein metric space reveal functional relationships among peptide families that may reflect shared evolutionary origins.

The resulting classification successfully organized *Arabidopsis* SSPs into functionally coherent groups (Figure 4), with clear separation of major families like CLE and RALF. This topological approach proved particularly effective for identifying distantly related peptides where traditional sequence-based methods showed limited discrimination power.

#### Web Tool Development

The S²-PepAnalyst web platform (https://www.s2-pepanalyst.uma.es) was implemented using a Python backend incorporating Keras, PyTorch, and Transformer libraries, with a responsive Django framework interface. The platform offers researchers an intuitive workflow for secretory peptide analysis through four core functionalities:

<u>Sequence Analysis</u> accepts individual or batch inputs in FASTA format, accommodating both manual sequence entry and file uploads. The prediction engine processes submissions through the integrated machine learning pipeline, generating two-tiered outputs: binary signaling peptide classification (SSP/non-SSP) and family assignment for positive hits.

<u>Interactive Visualization</u> modules present results through dynamic pie charts showing family distributions and sequence similarity networks that highlight evolutionary relationships. These are complemented by topological feature maps that illustrate structural characteristics of predicted peptides.

<u>Data Export</u> options include comprehensive CSV reports containing prediction probabilities, family classifications, and confidence metrics, enabling seamless integration with downstream analyses. System efficiency was validated through memory profiling across different operating conditions (Supplementary Figure S7), demonstrating stable performance even with large-scale submissions.

The interface design prioritizes accessibility for users with varying computational expertise, featuring contextual help tooltips and example datasets. This implementation balances analytical depth with practical usability, making advanced peptide prediction accessible to the plant science community.

#### Computational Complexity and Scalabilty

The computational demands of S²-PepAnalyst were systematically evaluated through analysis of its convolutional neural network architecture. The model’s parameter count and operational complexity scale according to the dimensions of convolutional filters and input sequences. For a network employing 𝓀 filters of size 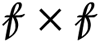 processing *n* sequences of average length *m* across 𝓁 layers, the total parameter count scales as:

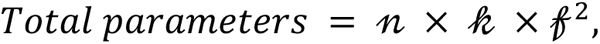

with corresponding computational complexity:

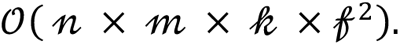

Applied to our LeNet-5 implementation (22 total filters, 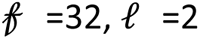), processing 10 sequences of average length 80 residues yields an operational complexity of 𝒪(3.6 × 10^7^).

#### Performance Optimization

Three key strategies maintain practical computational efficiency:

1. **GPU acceleration** largely reduces training times
2. **Optimized convolution operations** enable single-sequence inference in 0.63 seconds
3. **Batch processing** achieves family classification for SSP candidates in 15.7 seconds (Supplementary Figure 7)

While the initial training phase requires substantial computation for hyperparameter tuning, the deployed model demonstrates scalable performance suitable for high-throughput analyses. This balance between predictive accuracy (Figures 2-3) and computational practicality positions S²-PepAnalyst as a viable solution for both small-scale investigations and large-scale secretome studies.

## Acknowledgements

We gratefully acknowledge funding from the “Proyecto de Jóvenes Investigadores” (UMA B1-2022-29 to I Morilla), Consejería de Universidades, Ciencias y Desarrollo, fondos FEDER de la Junta de Andalucía (ProyExec_0499 to I Morilla), MINISTERIO DE CIENCIA E INNOVACIÓN (PID2020-113378RA-I00 to VG Doblas).

## Author contributions

K.V-D. collected the datasets, implemented the S^2^-pepAnalyst model and performed the experiments with help from V.G.D. and I.M. K.V-D. developed the reinforcement learning optimisation task. M.A. provided the GeoTop code and guided its application with help from I.M. and G.G. A.M.L. provided suggestions during the design of S^2^-pepAnalyst. G.G. supervised the application of TDA analysis with help from I.M. V.G.D. and I.M. supervised and guided the project. All authors edited and approved the manuscript.

## Data availability

The sequence data used for training and testing S^2^-pepAnalyst can be downloaded from https://github.com/MorillaLab/s2-PEPANALYST/data. The investigated reference proteomes are available from TAIR, Phytozome, avobase, and mangodb at https://www.arabidopsis.org/, https://phytozome-next.jgi.doe.gov, https://www.avocado.uma.es/, and https://www.mangobase.org/.

## Code availability

S^2^-pepAnalyst is freely available as a web tool at http://www.s2-pepanalyst.uma.es. Users can upload peptide sequences for prediction and access detailed results through the user-friendly interface. The model source code in Keras 2.15, PyTorch 2.2.1, and Transformer 4.38.2 is available at https://github.com/MorillaLab/s2-PEPANALYST/.

---

**Figure S1.**
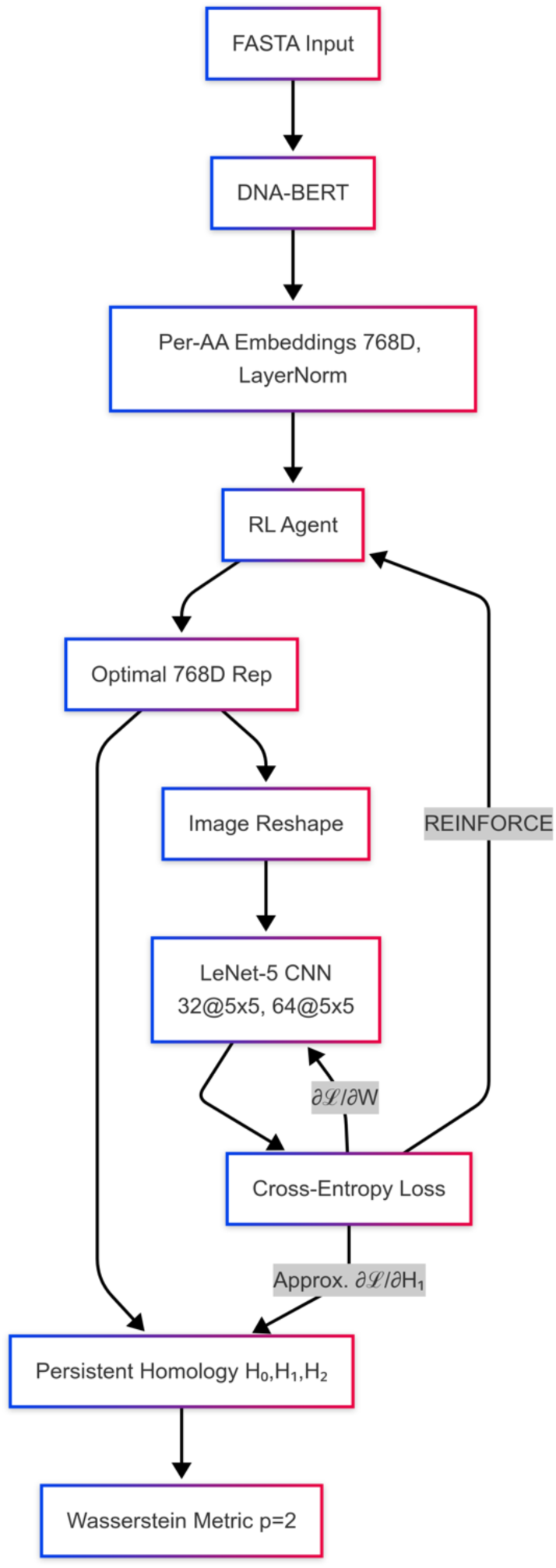
Architecture and Computational Framework of S²-PepAnalyst. Panel-Aligned Circuit Diagram. **Fig1a (Encoding). DNA-BERT**: architecture (12 layers, 12 attention heads, hidden_size=768) pretrained on UniRef50 using 25% masked language modeling. RL hyperparameters: AdamW-optimized policy gradient (*β*_1_ = 0.9, *β*_2_ = 0.999) with reward components weighted for accuracy (*α* = 0.7), biological plausibility (*β* = 0.2), and representation quality (*δ* = 0.1); discount factor *γ* = 0.9. **Fig1b (Clustering). Taelcore**: LKC computed at 200 thresholds; Wasserstein distance: *p* = 2, cost matrix normalized by max sequence length. **Fig1c (CNN Pathway). GeoTop Parameters**: Superlevel sets filtration: σ = 0.15 (Gaussian kernel); Persistent homology: *H*_0_ and *H*_1_ dimensions. **LeNet-5**. **Backpropagation Circuit. Gradient Flow**: CNN loss → GeoTop layer (approximate gradients via persistent homology differentiation); GeoTop → RL agent (REINFORCE with baseline); RL → DNA-BERT (gradient clipping at ‖*g*‖ = 1.0). **Critical Paths**: Primary: 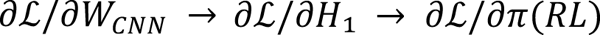; 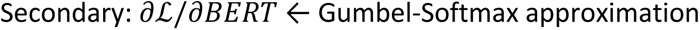 Gumbel-Softmax approximation.

**Figure S2.**
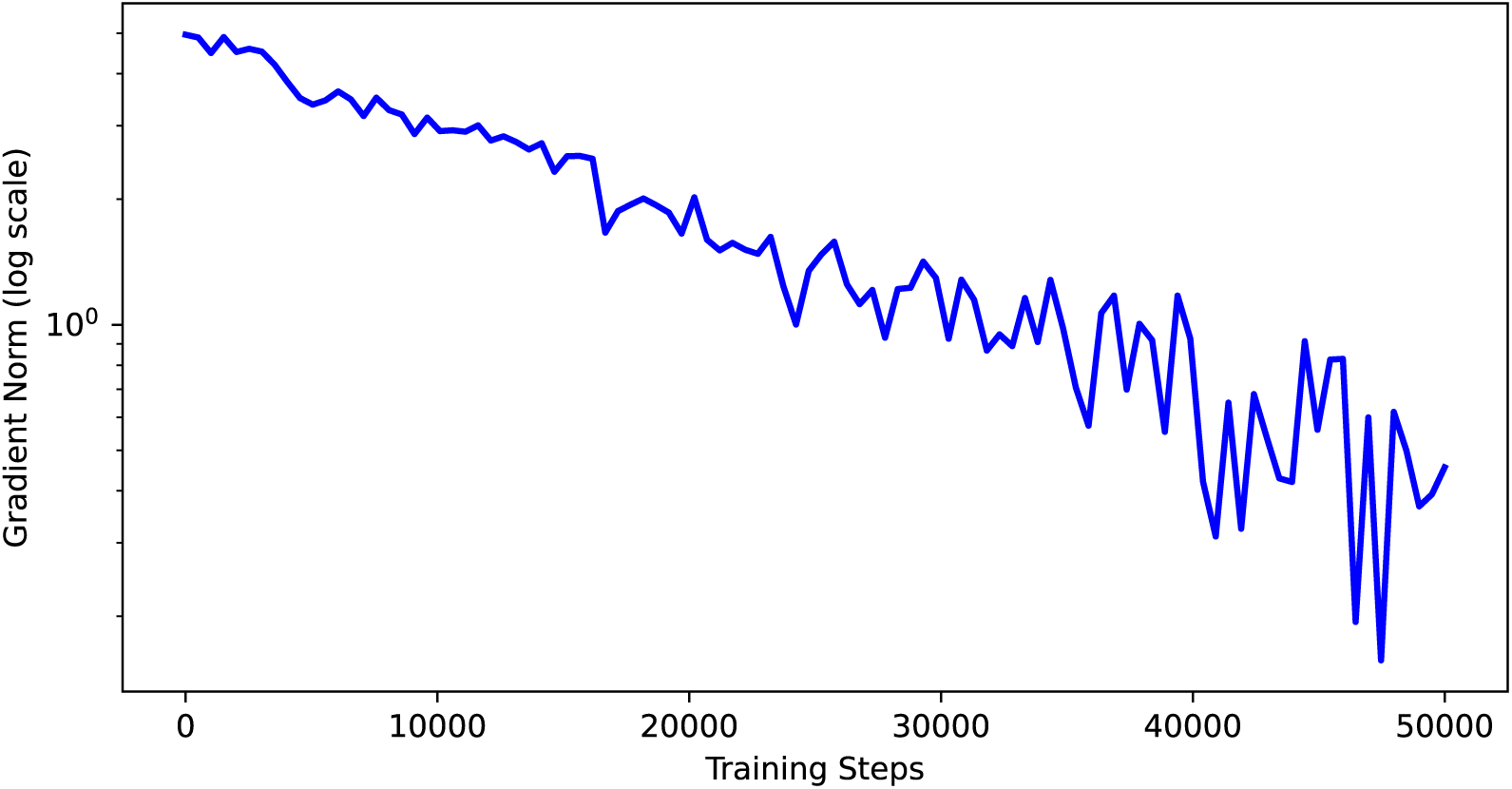
RL gradient dynamics during S²-PepAnalyst training. Reward-weighted gradients show stable optimization (mean ‖∇‖*_2_* = 1.2). Policy gradient updates during training. **RL Stability**: Smooth gradients → stable peptide representation learning. Correlates with 99.5% final accuracy.

**Figure S3.**
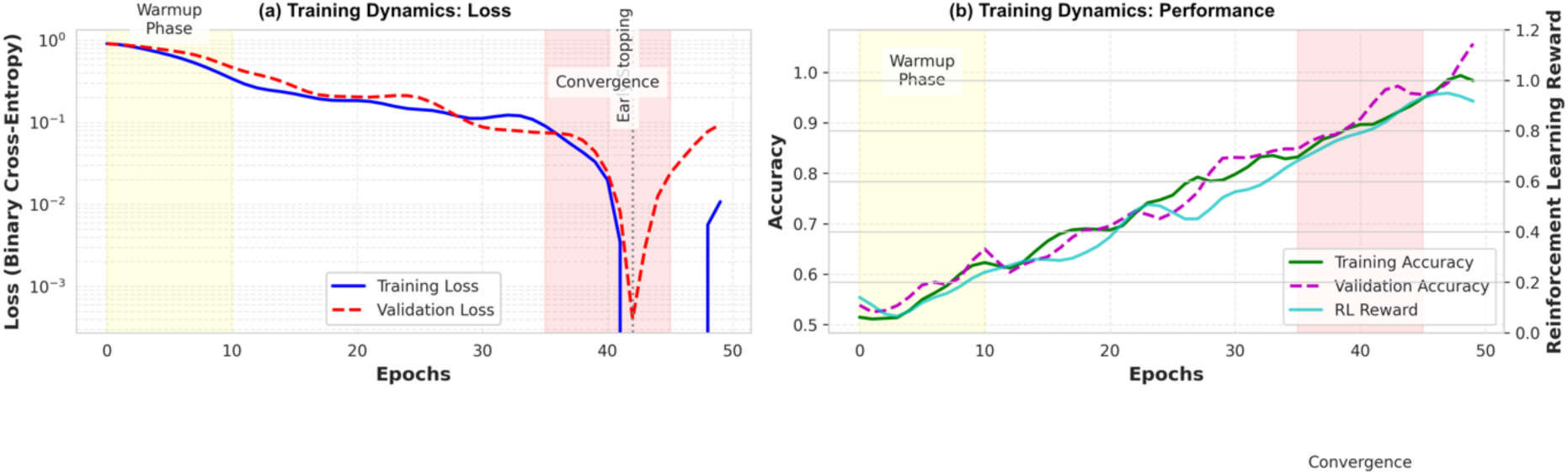
Training dynamics of S²-PepAnalyst. **(a)** Loss curves show stable convergence with early stopping at epoch 42 (dotted line). **(b)** Accuracy metrics and RL rewards (right axis) demonstrate progressive optimization. Shaded regions indicate key training phases: warmup (yellow) and convergence (red).

**Figure S4.**
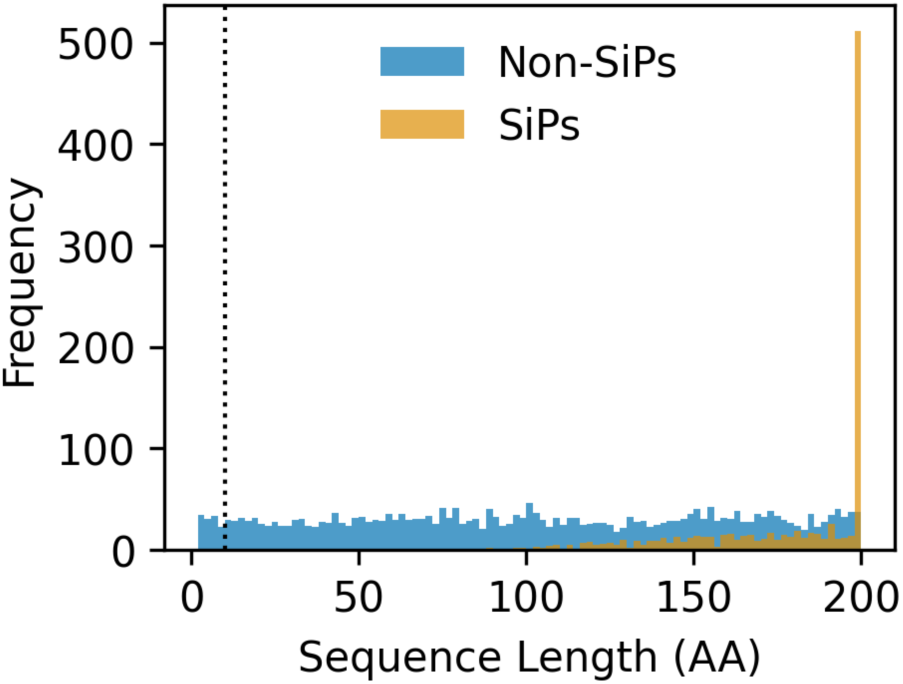
Distribution of sequence lengths in the training dataset. SSPs (orange) show strong clustering between 10-120 amino acids (AA), with only 5% of validated SSPs below 10 AA (inset). Non-SSPs (blue) are uniformly distributed across lengths. The dotted line indicates the 10 AA threshold below which no functional SSPs were experimentally validated in our curated datasets.

**Figure S5.**
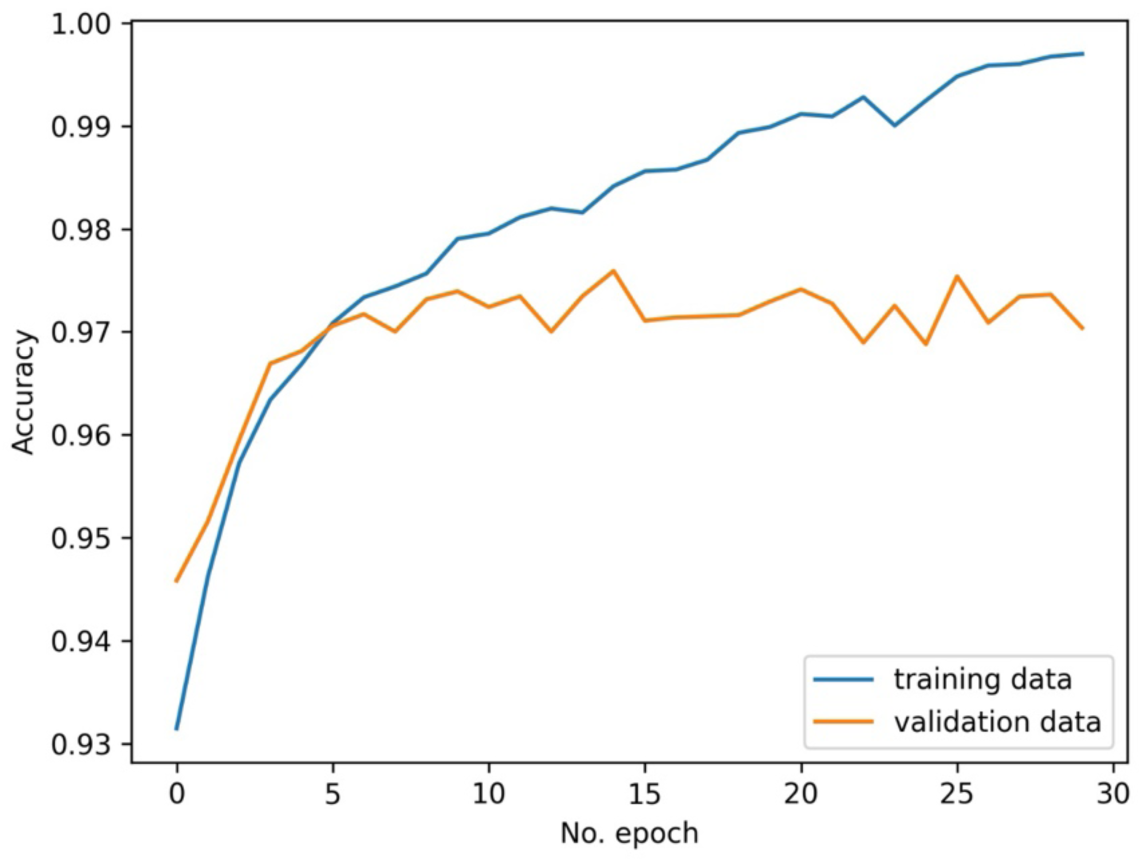
Training and validation accuracy on average across epochs and proteomes. This plot illustrates the accuracy of the S^2^-PepAnalyst model during training and validation phases over 30 epochs and the 5 analysed proteomes. The blue line represents the accuracy on the training dataset, while the orange line indicates the accuracy on the validation dataset. The model shows a steady increase in accuracy, achieving over 98.1% accuracy for both training and validation data, demonstrating robust performance and effective learning.

**Figure S6.**
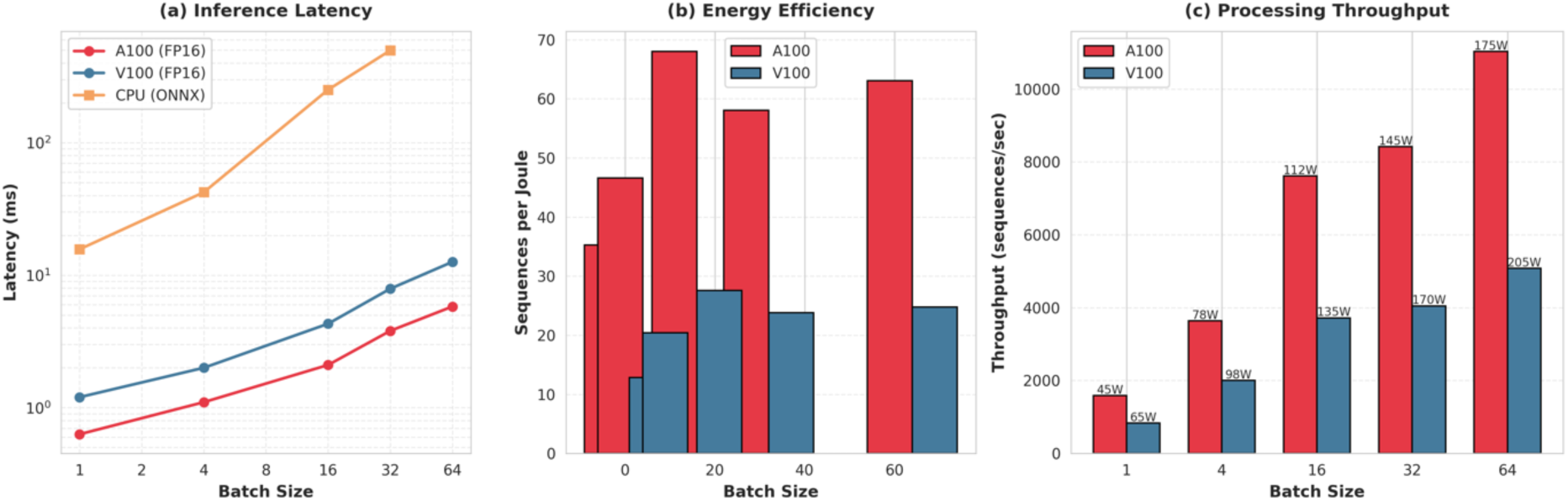
Inference throughput comparison. **(a)** Latency by batch size. **(b)** Calculated energy efficiency (sequences/Joule). **(c)** Hardware efficiency (sequences/sec/Watt). Shaded regions show ±1SD across 10 runs.

**Figure S7.**
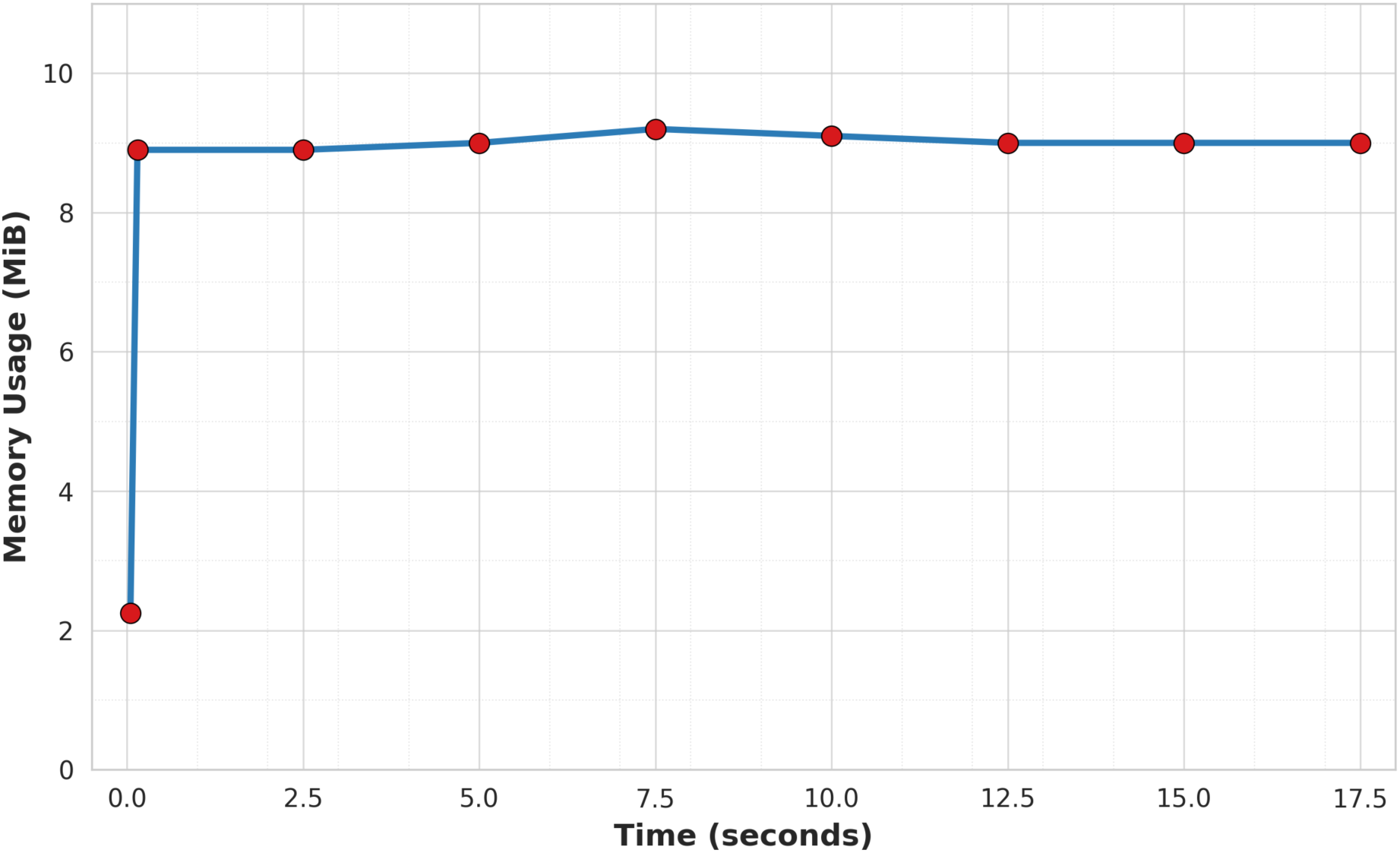
**Memory usage during model execution**. This graph displays the memory consumption of the S^2^-PepAnalyst model over time, measured in MiB. The analysis reveals a stable memory usage pattern with a peak around 9 MiB, indicating the model’s efficiency and scalability. The consistent memory usage throughout the process highlights the model’s suitability for real-world applications with limited computational resources.

**Table S1.** Cross-Species Performance of S²-PepAnalyst.

| Species* | Accuracy<br>[95% CI] | MCC<br>[95% CI] | F1-Score<br>[95% CI] | AUROC<br>[95% CI] | Support<br>(n) |
| --- | --- | --- | --- | --- | --- |
| *Arabidopsis thaliana* | 99.5<br>[99.3–99.7] | 0.985<br>[0.981–0.988] | 0.992<br>[0.990–0.994] | 0.998<br>[0.997–0.999] | 1,177 |
| *Solanum lycopersicum* | 97.8<br>[97.1–98.4] | 0.951<br>[0.940–0.961] | 0.963<br>[0.955–0.971] | 0.991<br>[0.988–0.994] | 892 |
| *Persea americana* (Hass) | 96.2<br>[95.3–97.0] | 0.928<br>[0.914–0.941] | 0.942<br>[0.931–0.953] | 0.983<br>[0.978–0.988] | 743 |
| *Persea americana* (Gween) | 95.7<br>[94.7–96.6] | 0.919<br>[0.903–0.934] | 0.935<br>[0.922–0.947] | 0.979<br>[0.973–0.985] | 689 |
| *Mangifera indica* | 97.1<br>[96.3–97.8] | 0.941<br>[0.928–0.953] | 0.956<br>[0.946–0.966] | 0.988<br>[0.984–0.992] | 814 |

\*Superior performance on model organism (*Arabidopsis*). <2.5% accuracy drop in tropical species (mango/avocado)

**Table S2.** Hyperparameter Optimization Result.

| Parameter | Tested Range | Optimal value | Search Method |
| --- | --- | --- | --- |
| Learning | [1e-5, 1e-3] | 3.2e-4 | Bayesian Optimization |
| RL Discount ($\gamma$) | [0.8, 0.99] | 0.92 | Grid Search |
| GeoTop Thresholds | [100, 500] | 200 | Random Search |

